# Cryo-EM structure of the RADAR supramolecular anti-phage defense complex

**DOI:** 10.1101/2022.08.17.504323

**Authors:** Brianna Duncan-Lowey, Nitzan Tal, Alex G. Johnson, Shaun Rawson, Megan L. Mayer, Shany Doron, Adi Millman, Sarah Melamed, Taya Fedorenko, Assaf Kacen, Gil Amitai, Rotem Sorek, Philip J. Kranzusch

## Abstract

RADAR is a two-protein bacterial defense system which was reported to defend against phage by ‘editing’ messenger RNA. Here we determine cryo-EM structures of the RADAR defense complex, revealing RdrA as a heptameric, two-layered AAA+ ATPase and RdrB as a dodecameric, hollow complex with twelve surface-exposed deaminase active sites. RdrA and RdrB join to form a giant assembly up to 10 MDa, with RdrA docked as a funnel over the RdrB active site. Surprisingly, our structures reveal a RdrB active site that targets mononucleotides, not RNA. We show that RdrB catalyzes ATP-to-ITP conversion in vitro and induces the accumulation of inosine mononucleotides during phage infection in vivo, limiting phage replication. Our results define ATP mononucleotide deamination as a determinant of RADAR immunity and reveal supramolecular assembly of a nucleotide-modifying machine as a novel mechanism of anti-phage defense.

## Introduction

Bacteria encode a rich and highly diverse arsenal of anti-phage immune systems, allowing them to mitigate phage infections (Bernheim and Sorek, 2020; Hampton et al., 2020). While most bacteria carry at least one restriction enzyme and about half of them encode CRISPR-Cas as part of their immune arsenal, other defense systems are sparsely distributed in microbial genomes (Payne et al., 2021; Tesson et al., 2022). Over sixty anti-phage defense systems have been discovered in the past few years (Doron et al., 2018; Gao et al., 2020; LeRoux and Laub, 2022; Millman et al., 2022; Rousset et al., 2022) and while the mechanisms of action of a minority of them have been determined (Bernheim et al., 2021; Garb et al., 2021; Johnson et al., 2022; LeRoux et al., 2022; Millman et al., 2020; Tal et al., 2022; Whiteley et al., 2019), the vast majority remain unexplored. It is estimated that an average bacterial genome contains more than five defense systems, with some bacteria encoding as many as 57 such systems (Tesson et al., 2022).

An intriguing defense system that was recently discovered is called RADAR (“Restriction by an Adenosine Deaminase Acting on RNA”) (Gao et al., 2020). The RADAR system comprises two genes, *rdrA* that encodes a protein with an ATPase domain, and *rdrB*, encoding a predicted adenosine deaminase domain protein (Figure 1A). Both RdrA and RdrB are large proteins, with an average size of ~900 amino acids each. The RADAR operon from *Citrobacter rodentium* DBS100 was heterologously expressed in *E. coli* and shown to confer defense against phages T2, T3, T4 and T5, in a manner dependent on both RdrA and RdrB (Gao et al., 2020). The RADAR system was proposed to confer defense by editing of adenosine to inosine residues in RNA molecules of the bacterial host, thus leading to abortive infection via growth arrest or death of the infected cells (Gao et al., 2020).

**Figure 1.**
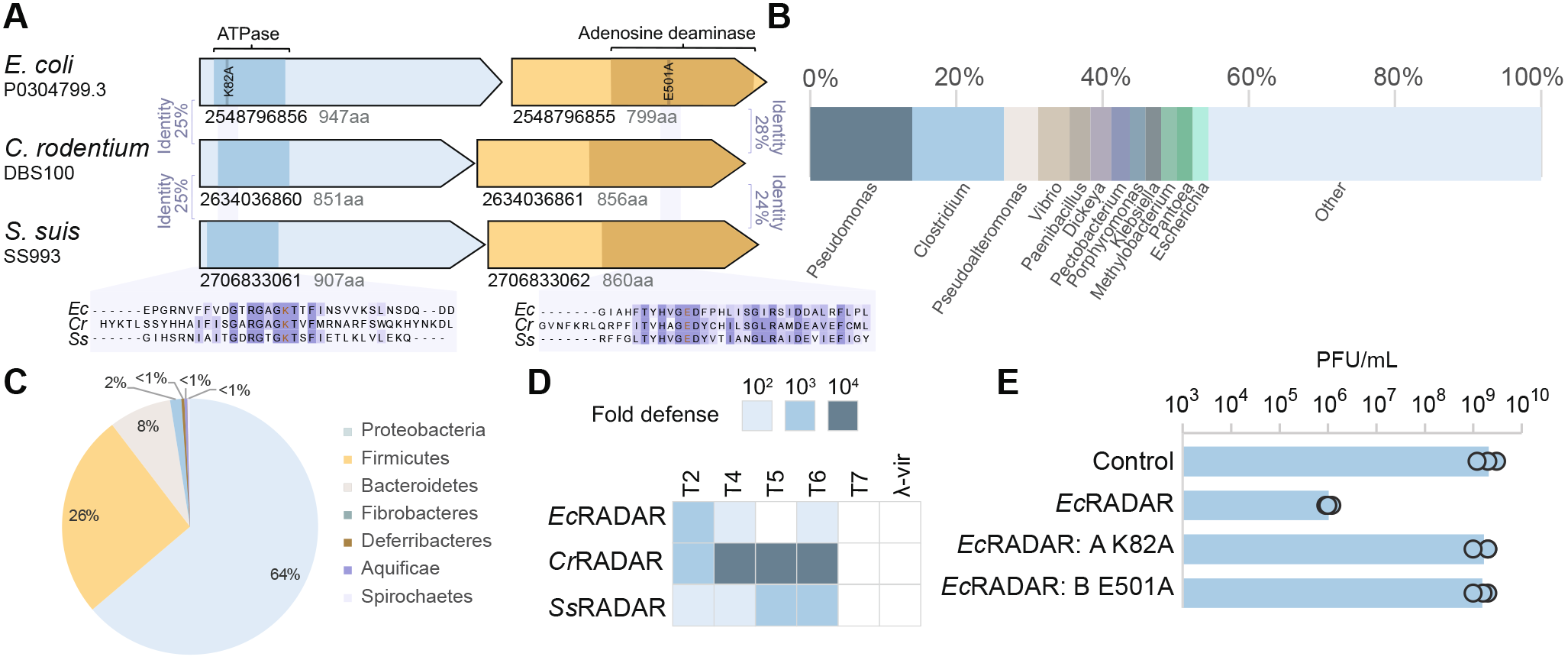
Diverse RADAR systems protect E. coli from phage replication. (A) RADAR systems studied here. (B) Genera of bacteria encoding RADAR. (C) Phylum distribution of RADAR-encoding bacteria. (D) RADAR systems defend against phages. RADAR systems were cloned into plasmids and transformed into E. coli MG1655. Fold defense was measured using serial dilution plaque assays. Data represent an average of three replicates (see detailed data in Figure S1, S2). (E) Effect of point mutations on the defensive activity of EcRADAR. Data represent plaque-forming units per mL (PFU mL−1) of T2 phage infecting control cells, EcRADAR-expressing cells, and two strains mutated in the predicted ATPase or deaminase domains. Shown is the average of three replicates, with individual data points overlaid.

In this study we used cryo-electron microscopy (cryo-EM) to determine the molecular structure of the RADAR defense complex. We find that RdrA forms a heptameric complex with a conserved AAA+ core and a previously uncharacterized C-terminal domain. RdrB, the adenosine deaminase protein, forms a highly atypical dodecameric shell that cages an enzymatic active site that has surprising homology to mononucleotide deaminases, not RNA-targeting enzymes. We demonstrate that the RdrA heptamer stably interacts with the RdrB dodecamer and forms a channel that feeds directly into the deaminase active site of RdrB. Multiple RdrA complexes can occupy the RdrB dodecamer, and these can together form a maximal supramolecular complex of up to 10 MDa. In contrast to previous hypotheses, we do not find strong evidence for RNA editing by the RADAR complex. Rather, we demonstrate that RADAR modifies the mononucleotides ATP and deoxy-ATP (dATP) to inosine triphosphate (ITP) and deoxy-ITP (dITP), respectively, in response to phage infection. This mononucleotide modification, which we observe both in vitro and in vivo, is suggested to block phage replication.

## Results

### Diverse RADAR systems protect *E. coli* from phage replication

A recent cross-genome defense system annotation effort detected 103 RADAR systems in ~22,000 analyzed genomes (Tesson et al., 2022). In the current study we analyzed a larger set of ~38,000 bacterial and archaeal genomes, and detected RADAR in 270 of these, confirming that this defense system is rare and occurs in <1% of analyzed sequenced genomes (Table S1). Despite its rarity, the RADAR system was widely distributed phylogenetically, occurring in 75 distinct genera spanning 7 phyla in our set (Figure 1B,C).

We selected the RADAR systems of *Escherichia coli* P0304799.3 and *Streptococcus suis* SS993 for further experimental investigation, as well as the previously studied *C. rodentium* DBS100 RADAR (Gao et al., 2020). Proteins in these systems vary substantially in amino acid sequence: it is only possible to align about one quarter of RdrA, largely spanning the ATPase domain, between pairs of RADAR systems in our set, and even in this aligned region, only 22–25% of the residues are identical (Figure 1A). Similarly, alignable regions between RdrB proteins span only the C-terminal portion which harbors part of the adenosine deaminase domain, with 24–28% sequence identities in aligned regions (Figure 1A). Such an extensive divergence in protein sequences is typical to proteins involved in immunity (Daugherty and Malik, 2012) and has been observed in other rare defense proteins such as the prokaryotic viperin and gasdermin anti-phage defense operons (Bernheim et al., 2021; Johnson and Wein et al., 2022). Despite the divergence in sequences, the predicted active sites were conserved in both RdrA and RdrB (Figure 1A).

When heterologously expressed in *E. coli* MG1655, all RADAR systems conferred defense against the closely related T-even phages T2, T4 and T6, with two of them also defending against T5, confirming the previous report on *C. rodentium* RADAR (Figure 1D; Figure S1) (Gao et al., 2020). Mutations in the *E. coli* RADAR, predicted to inactivate the ATPase site of RdrA or the deaminase active site of RdrB abolished defense, verifying, also in agreement with previous reports (Gao et al., 2020), that both enzymatic functions are necessary for defense (Figure 1E).

### Structure of RdrA reveals a two-layered heptameric ATPase assembly with a unique C-terminal domain

To define the role of RdrA in phage defense, we used cryo-EM to determine a 2.5 Å structure of *E. coli* RdrA (Table S2, Figure S2A,B,C). The cryo-EM structure of RdrA reveals a heptameric assembly with two layers of interlocking protein domains (Figure 2A). The top layer of the RdrA assembly contains seven N-terminal AAA+ ATPase domains with an active site formed between each pair of neighboring subunits. Extending from the N-terminal AAA+ domain, the RdrA C-terminus is a largely alpha-helical lobe that interlocks around the assembly to form a flared bottom-layer of the heptameric complex. We analyzed minor complexes in the *E. coli* RdrA data and determined a series of additional structures including a 2.5 Å structure of RdrA with a single break between subunits and a 2.4 Å structure of RdrA with breaks between two subunits, suggesting conformational flexibility within the heptameric ring-like assembly (Figure S2D,E; Table S2).

**Figure 2.**
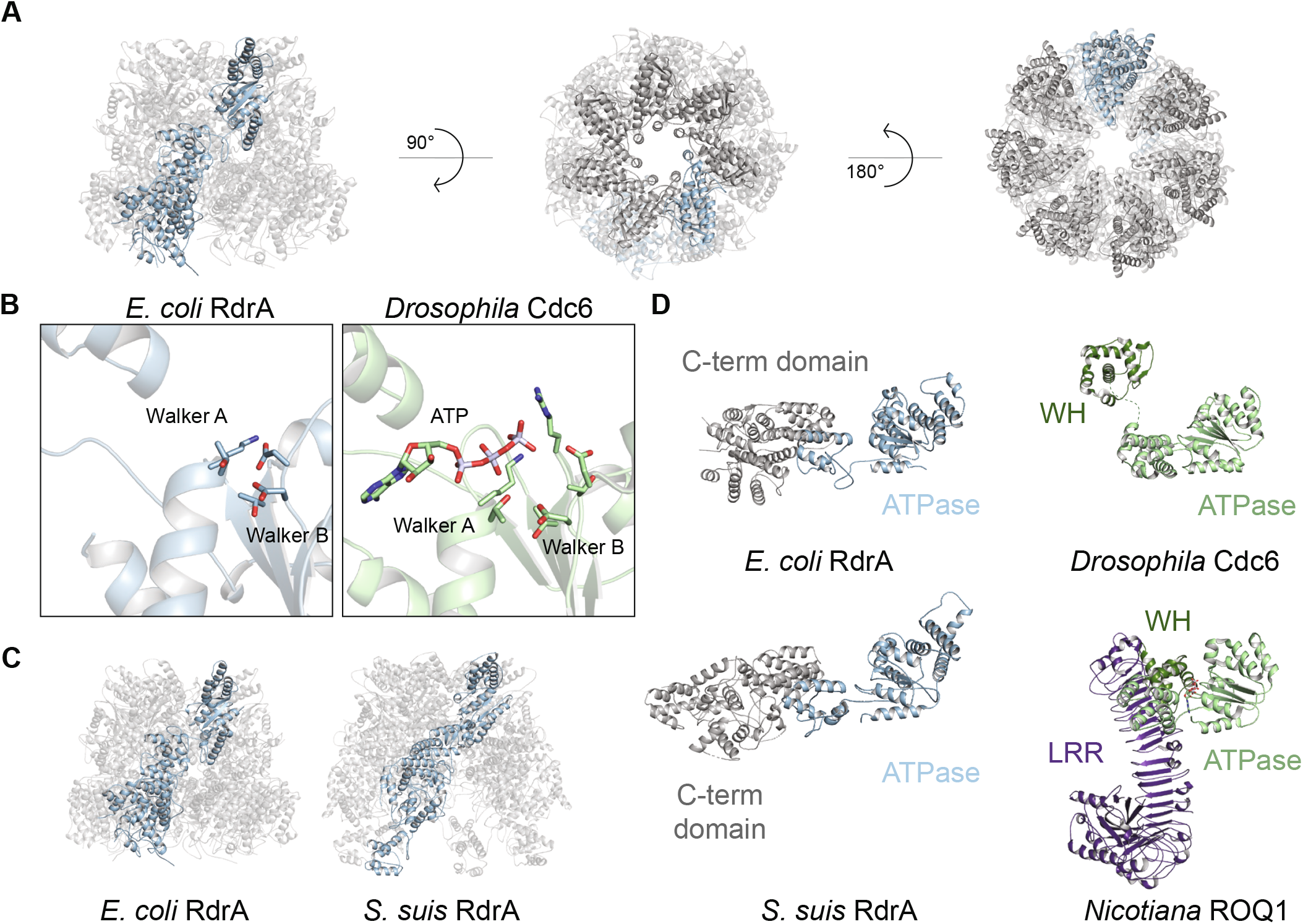
Structure of RdrA reveals a two-layered heptameric ATPase assembly with a unique C-terminal domain. (A) Cartoon representation of E. coli RdrA cryo-EM structure (B) Comparison of E. coli RdrA and Drosophila Cdc6 active sites, containing the well-conserved Walker A and Walker B motifs. (C) Comparison of E. coli RdrA and S. suis RdrA, highlighting the flipping out of the C-terminal helical bundle at the base of the protein. (D) Comparison of E. coli and S. suis RdrA to ATPase domain-containing proteins: Drosophila Cdc6 (PDB 7JGR), part of the origin recognition complex, and Nicotiana ROQ1 (PDB 7JLU, 7JLV), a plant NLR that sense the pathogen effector protein XopQ.

AAA+ ATPase domain-containing proteins comprise a diverse family of molecular machines that hydrolyze ATP to rearrange target binding partners and mechanically funnel polypeptide and nucleic acid substrates (Seraphim and Houry, 2020). Similar to canonical AAA+ proteins like *Drosophila* Cdc6 (Schmidt and Bleichert, 2020), the RdrA AAA+ ATPase domain contains highly conserved features associated with enzymatic activity including Walker A (G81, K82, T83) and Walker B (D221, D222) motifs, the sensor 1 motif (D253), and a sensor 2 motif (R377) positioned to catalyze ATP hydrolysis (Figure 2B, Figure S2F). Although RdrA contains the core elements associated with enzymatic activity, the RdrA assembly exhibits several distinguishing features that are distinct compared to previously defined structural clades of AAA+ ATPase proteins (Erzberger and Berger, 2006). In contrast to most closed-ring AAA+ proteins that form hexameric assemblies, RdrA is an atypical complex with 7 repeating subunits. RdrA also lacks a pre-sensor 1 beta-hairpin insertion common in many clades of AAA+ proteins (Miller and Enemark, 2016; Seraphim and Houry, 2020), and instead contains two large insertions not associated with designated AAA+ clades. The first RdrA insertion occurs after strands β2 and β3 where a canonical single helix α2 is replaced with a four-helix bundle that forms a crown at the top of the heptameric complex (discussed further in Figure 4). The second RdrA insertion occurs after the sensor 1 motif and includes three helices between strands β4 and β5 that reach into the central channel of the heptamer (Figure S2F).

In contrast to the N-terminal AAA+ domain, the alpha-helical RdrA C-terminus contains no detectable homology with other known structures. To better understand the RdrA C-terminal domain, we next determined a 2.5 Å cryo-EM structure of the distantly related (< 20% amino acid identity) RdrA from *Streptococcus suis* (Figure 2C, Figure S2 G,H, Table S2). *S. suis* RdrA adopts a similar two-layered heptameric architecture confirming conservation of the C-terminal alpha-helical lobe despite high sequence variability. Interestingly, *S. suis* RdrA particles existed primarily as 14-mer complexes with the RdrA stacked head-to-tail (Figure S2G). In this conformation, the final four alpha helices of the RdrA C-terminus undergo a dramatic ~120° reorientation relative to the *E. coli* RdrA structure and flip out to form a protein-protein binding interface (Figure 2C). A notable immune protein that shares an AAA+ ATPase core structurally homologous to the RdrA ATPase is the plant NOD-like receptor ROQ1 which senses *Xanthomonas* pathogens (Figure 2D). In ROQ1-dependent immunity, sensing requires direct protein-protein interaction between the ROQ1 C-terminal leucine rich repeat domain and a *Xanthomoas* effector protein, which then induces downstream immune signaling (Martin et al., 2020). The C-terminal alpha-helical domain of RdrA may similarly act as the pathogen-sensing domain in response to an unknown phage signal.

### Structure of RdrB reveals an adenosine deaminase dodecamer

The second RADAR defense protein, RdrB, is an ~90 kDa protein predicted to encode an adenosine deaminase domain (Figure 1A) (Gao et al., 2020). Cryo-EM analysis of *E. coli* RdrB showed an assembly of a rigid, higher-order complex, and we determined a structure to 2.1 Å (Table S2, Figure S3A–C). *E. coli* RdrB forms a dodecameric assembly with 12 RdrB protomers joined to create a large hollow shell that is ~160 Å in diameter (Figure 3A,B).

**Figure 3.**
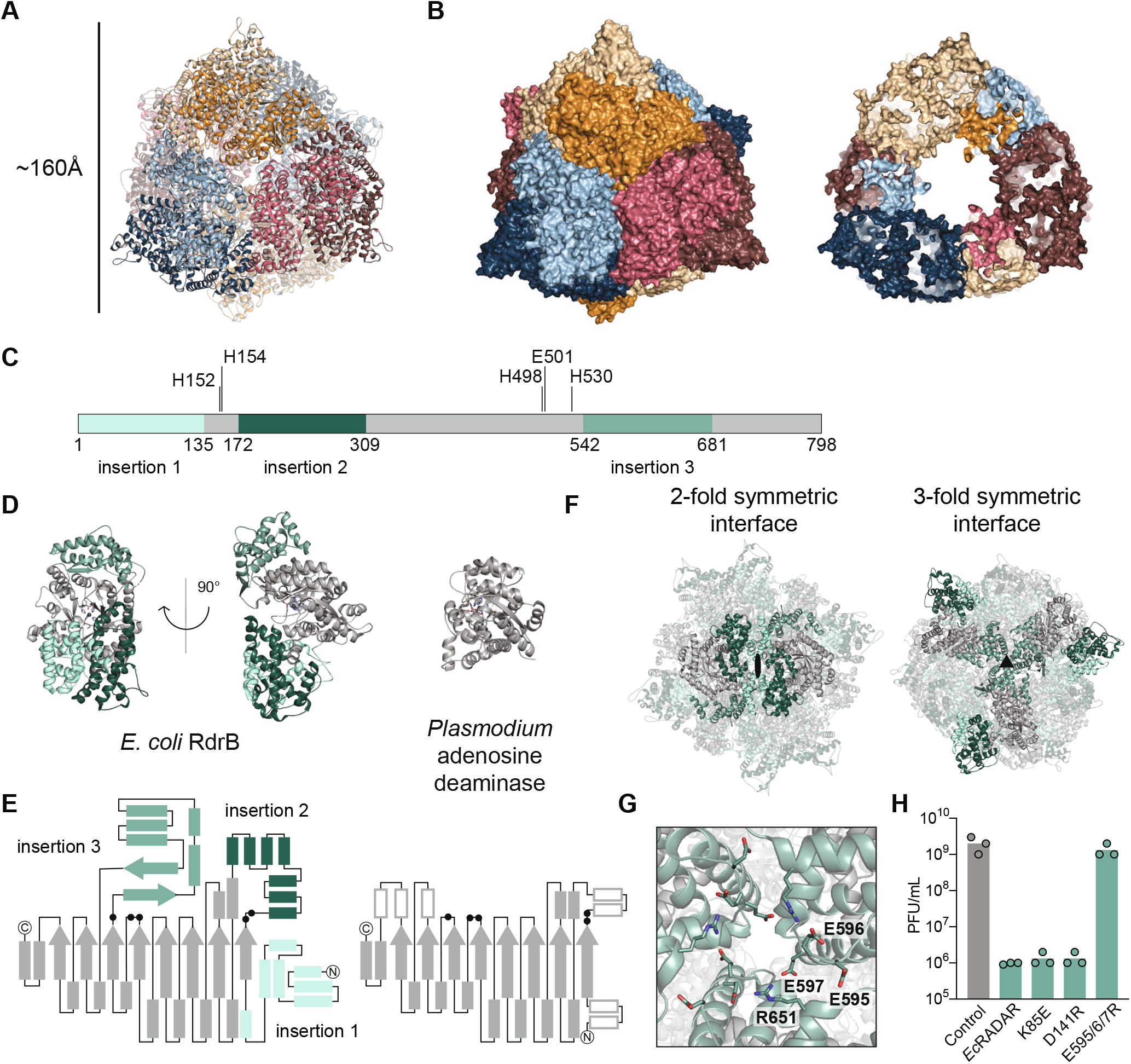
Structure of RdrB reveals an adenosine deaminase dodecamer. (A) Cartoon representation of E. coli RdrB dodecamer, with each protomer in a different color. (B) Surface representation of E. coli RdrB dodecamer (left), and slice through showing hollow center (right) (C) Schematic representation showing insertions in RdrB relative to other adenosine deaminases. (D, E) Structure (D) and schematic (E) representation of single RdrB monomer highlighting insertions in shades of green (left) compared to Plasmodium adenosine deaminase (PDB 2PGF) (right). Active site residues are shown as black dots in the schematic (E). (F) Insertions 1 and 2 form two-fold symmetric dodecamer interface and insertion 3 forms the three-fold symmetric dodecamer interface. (G) Close up view of 3-fold interface shown in (F) highlighting key residues, including those mutated to disrupt the interface in (H). (H) Effect of point mutations suggested to disrupt the RdrB multimerization interfaces on the defensive activity of EcRADAR. Data represent plaque-forming units per mL (PFU mL−1) of T2 phage infecting control cells, EcRADAR-expressing cells. Shown is the average of three replicates, with individual data points overlaid. The same control replicates are shown in Figure 1E and 4G for reference.

**Figure 4.**
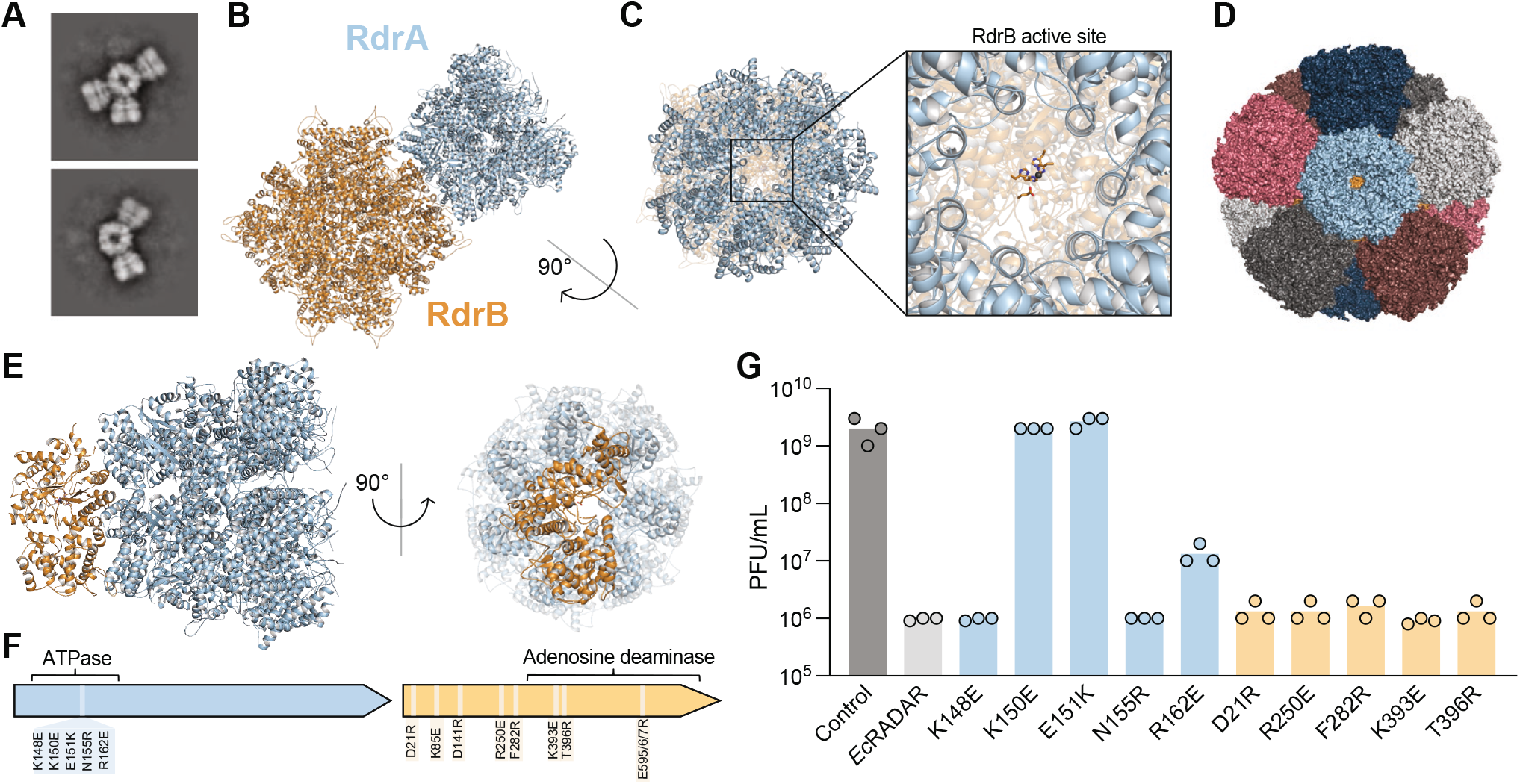
Rdr and RdrB form a supramolecular complex required for anti-phage defense. (A) 2D class averages of cryo-EM RdrA-RdrB mixture showing many arrangements with different numbers of RdrA “petals” surrounding a RdrB core. (B) Cartoon representation of RdrA heptamer (Blue) in complex with a RdrB dodecamer (yellow). (C) View down the central channel of RdrA heptamer (blue) showing access to RdrB active site directly at the base (yellow). (D) Model of RdrB dodecamer with a RdrA heptamer bound to each of the 12 protomers, showing there is space to accommodate each RdrA heptamer in this hypothetical “saturated” complex. (E) RdrB monomer and RdrA heptamer, colored as in (D), showing the footprint of RdrA is largely restricted to a single RdrB protomer. (F) Location of mutation in RdrA and RdrB. (G) Effect of point mutations suggested to disrupt RdrA-RdrB complex on the defensive activity of EcRADAR. Data represent plaque-forming units per mL (PFU mL−1) of T2 phage infecting control cells. Shown is the average of three replicates, with individual data points overlaid. The same control replicates are shown in Figure 1E and 3G for reference.

Structural analysis of the *E. coli* RdrB complex reveals three large insertions to the adenosine deaminase core that mediate higher order dodecameric assembly. Compared to a typical adenosine deaminase domain like *Plasmodium vivax* ADA (Larson et al., 2008), RdrB contains an N-terminal extension “insertion 1”, a unique “insertion 2” that divides the catalytic core, and an “insertion 3” after the final catalytic histidine H530 (Figure 3C–E). Insertions 1 and 2 form a 2-fold symmetrical interface that joins adjacent adenosine deaminase core domains (Figure 3F). Insertion 3 forms a hydrophilic interface consisting of alternating glutamate and arginine residues that create a 3-fold symmetry axis and enable assembly of the large dodecameric shell (Figure 3F, G). Some adenosine deaminases are known to function as dimers (Figure S3D) (Zavialov et al., 2010), but to our knowledge assembly of a large, shelled complex is a feature unique to RADAR RdrB. We made several mutations along the interfaces between RdrB protomers (Figure 3H); charge disrupting mutations to the conserved glutamate residues that line interface 3 (E595R, E596R, E597R) result in complete loss of anti-phage defense and demonstrate that RdrB complex assembly is essential for RADAR function (Figure 3G,H).

### RdrA and RdrB form a supramolecular complex required for anti-phage defense

RdrA and RrdB are encoded in tandem within all known RADAR systems and each protein is essential for anti-phage defense (Figure 1) (Gao et al., 2020). We therefore hypothesized that the two protein assemblies may directly interact. We analyzed mixed samples of purified *E. coli* RdrA and RdrB by negative-stain EM and observed gigantic flower-like arrangements with a central hub surrounded by up to six “petals” (Figure S4A). Further analysis with cryo-EM confirmed specific RdrA-RdrB co-complex formation (Figure 4A) and allowed us to determine a 6.7 Å structure of a representative single-petal complex (Figure S4B–E, Table S2). Placement of the high-resolution *E. coli* RADAR protein models in the RdrA-RdrB cryo-EM map reveals that the RdrB dodecamer forms the central hub and RdrA heptamers dock on the outside to create the extended petals observed by negative-stain EM (Figure 4B).

In the RdrA-RdrB complex, RdrA is positioned such that the central heptameric channel funnels directly into the catalytic core of RdrB (Figure 4C). In agreement with negative-stain EM analysis of multi-petal complexes, the RdrA heptamer footprint sits primarily over a single RdrB protomer and leaves room for a total of 12 docked RdrA complexes on the RdrB core dodecameric shell (Figure 4D,E). Nearly all contacts between RdrA and RdrB are mediated by residues that reside in an RdrA alpha-helical insertion that forms a crown on top of the AAA+ ATPase domain (Figure 2; Figure S4F). Modeling of a complete ~10 MDa flower-like RdrA-RdrB complex demonstrates that the RdrB adenosine deaminase active site remains solvent-accessible, suggesting that supramolecular assembly could represent that active state of the RADAR defense complex. We introduced several mutations predicted to disrupt the formation of the RdrA-RdrB complex, including charge-swap mutations of conserved residues in the RdrA crown (K150E, E151K, R162E) (Figure 4F, S4F), and observed that mutations to residues involved in RdrA-RdrB complex formation significantly decrease or ablate RADAR defense (Figure 4G). Together, these results reveal that supramolecular complex formation is an essential step of RADAR anti-phage defense.

### Structural analysis of RdrB suggests targeting of nucleotide substrates

It was previously suggested that the RADAR system from *C. rodentium* edits RNAs of the host bacteria to promote abortive infection (Gao et al., 2020). To test whether the *E. coli* RADAR edits host RNA during infection, we performed whole-transcriptome RNA sequencing of cells infected by T2 or T4 phages, at multiplicity of infection of 2, at 15, 27 and 120 min from the time of infection. Sequenced RNA reads with RNA-edited adenosines (A) should show A-to-G mismatches when aligned to the reference genome(Suzuki et al., 2015). However, regardless of the time point examined, the vast majority of adenosine residues (>99%) in expressed RNAs remained adenosines and were not converted, neither in the bacterial nor the phage transcriptome (Figure 5A; Figure S5A; Table S3). We repeated the experiment with cells expressing the RADAR from *C. rodentium*, infected by phage T2, but did not see strong signatures of A-to-G RNA editing (Figure S5A; Table S3). These results do not support the hypothesis that RNA is rampantly edited by the RADAR system.

**Figure 5.**
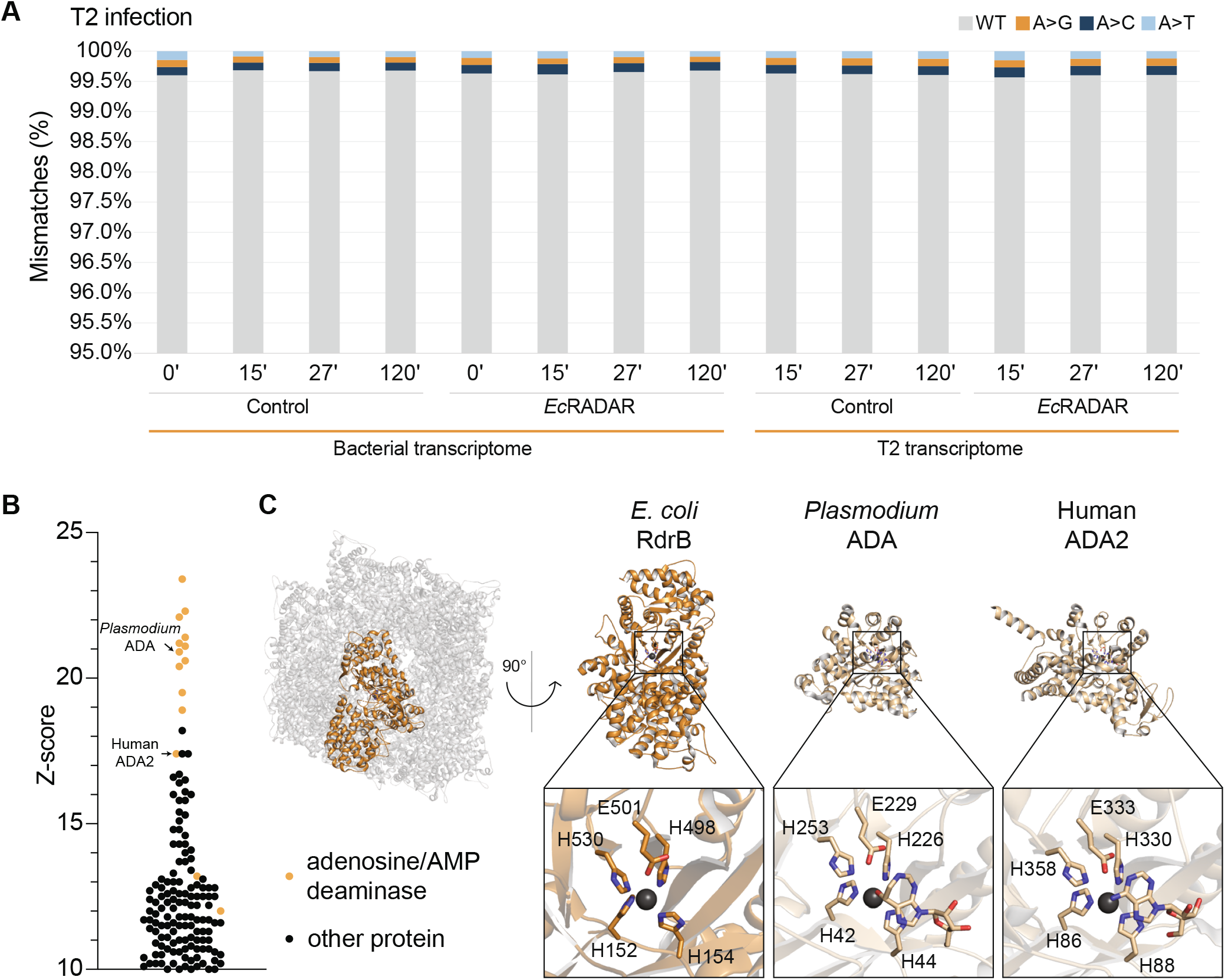
Structural analysis of RdrB suggests targeting of nucleotide substrates. (A) Mismatches between sequenced RNA and genomic DNA in adenine positions. RNA, extracted from EcRADAR-expressing or control cells infected by phage T2, was sequenced and the resulting reads were aligned to the reference sequences of the E. coli host and the T2 phage genomes. Shown in the rate of mismatches in all expressed adenosine positions mapped to non-rRNA genes. X-axis depicts the time from the onset of infection, with t=0 reflecting uninfected cells; y-axis is the observed rate of mismatches. (B) DALI Z-score of protein structures similar to RdrB, showing homology to adenosine and AMP deaminases (highlighted in yellow) (C) Cartoon representation of RdrB dodecamer, highlighting a single monomer in yellow (left), and comparison to other adenosine deaminase active sites (right).

We next considered the possibility that RADAR edits specific adenine residues in the bacterial transcriptome rather than consistently throughout the transcriptome. It was previously shown that RADAR from *C. rodentium* converts adenines to inosines specifically in RNA loops that reside in stem-loop secondary structures of highly expressed bacterial RNAs (Gao et al., 2020). We examined the 49 RNA loops reported by Gao *et al.* as heavily edited by RADAR during T2 infection but could not find evidence for substantial editing of these positions in the *E. coli* or the *C. rodentium* RADAR RNA-seq data (Figure S5A; Table S3).

As robust RNA editing was not observed *in vivo* during RADAR anti-phage defense, we analyzed the RdrB structure to assess potential targets for deaminase activity. Comparison of *E. coli* RdrB against all structures in the Protein Data Bank revealed that *E. coli* RdrB has no structural similarity to Adenosine deaminases acting on RNA (ADAR) proteins, which are part of the cytidine deaminase superfamily of enzymes that use a mechanistically distinct active site to edit RNA (Figure 5B, Figure S5B) (Goodman et al., 2012; Holm, 2022). Instead, all the top hits with structural similarity to *E. coli* RdrB (Z-score > 20) are adenosine or AMP deaminases that deaminate the adenosine within unphosphorylated or monophosphorylated monomeric substrates, as opposed to within RNA (Figure 5B). Importantly, *E. coli* RdrB shares the same active site architecture of a histidine tetrad that coordinates a zinc ion (*E. coli* RdrB H152, H154, H498, H530) and a catalytic glutamate residue (*E. coli* RdrB E501) as known adenosine deaminases like *Plasmodium vivax* ADA (PDB: 2PGF) and the human protein ADA2 (PDB: 3LGG) (Figure 5C) (Larson et al., 2008; Zavialov et al., 2010). This structural analysis suggests that RdrB enzymatic activity may target mononucleotide substates.

### ATP-to-ITP and dATP-to-dITP conversion mediates RADAR anti-phage defense

As the structural analysis of RdrB pointed to homology to adenosine deaminases acting on mononucleotides, we examined the possibility that RADAR converts ATP or deoxy-ATP to inosine derivatives during phage infection. We infected *E. coli* RADAR-expressing cells with phage T4 and collected cell lysates from several time points during infection. Analysis of filtered cell lysates via mass spectrometry showed accumulation of both ITP and dITP during infection, as early as 5 min from the onset of infection (Figure 6A). Similar experiments with cells in which RdrA or RdrB were inactivated by point mutations in their active sites showed no ITP or dITP accumulation, suggesting that the specific activity of these two proteins is necessary for mononucleotide conversion (Figure 6A). Accumulation of ITP and dITP was also observed in cells expressing RADAR from *C. rodentium* during infection by phage T2 (Figure S6A). These results show that RADAR generates both ITP and dITP in response to phage infection.

**Figure 6.**
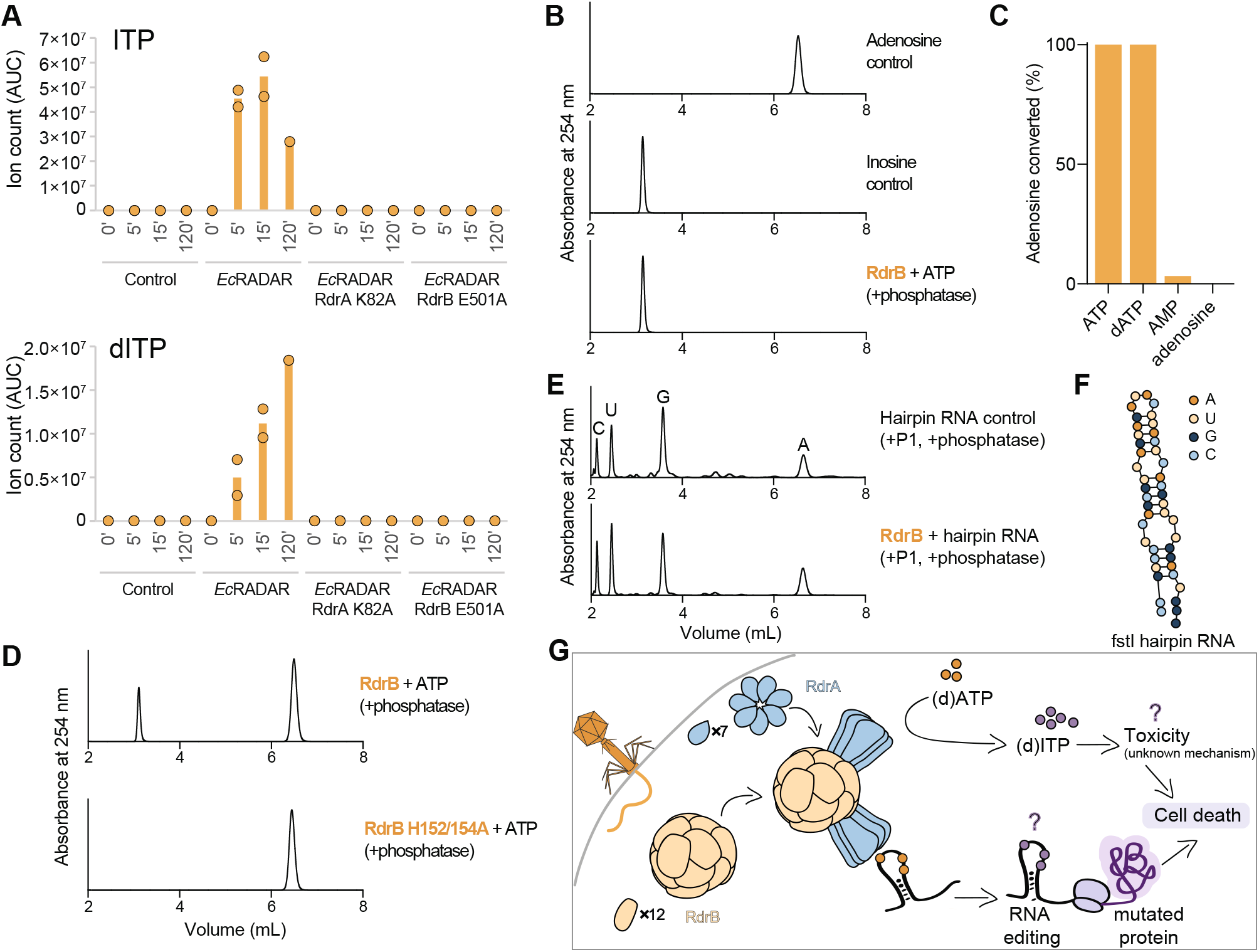
RADAR mediates ATP-to-ITP conversion in anti-phage defense. (A) Ion count (area under curve) of ITP or dITP (respectively) in lysates extracted from WT EcRADAR containing cells, as well as RADAR mutated in RdrA-K82A or RdrB-E501A, as measured by LC-MS/MS. The x-axis represents min after infection, with zero representing non-infected cells. Cells were infected by phage T4 at an MOI of 2 at 37°C. Bar graphs represent the average of two biological replicates, with individual data points overlaid. (B) HPLC analysis of ATP after incubation with purified E. coli RdrB, showing deamination of ATP to ITP. (C) Summary of HPLC analyses of 1 mM different nucleotide substrates incubated with 1 μM RdrB for 30 minutes, demonstrating robust deamination of tri-phosphorylated substrates, but not monophosphorylated or unphosphorylated substrate. Full HPLC traces shown in Figure S6B. (D) Mutation of conserved histidine residues within the active site of RdrB are required for deaminase activity, and RdrB H152/154A mutant no longer converts ATP to ITP. (ED) HPLC analysis of nucleotides after digestion of hairpin RNA to assess the identity of each base. Incubation of hairpin RNA with RdrB, in conditions where ATP is robustly converted to ITP (B), does not lead to deamination of adenosine within the RNA. (F) Schematic of fstI hairpin RNA used in (E). Hairpin was digested after incubation with RdrB, and mononucleotides were visualized by HPLC. (G) A proposed schematic for the mechanism of action of the RADAR defense system.

Given the accumulation of ITP and dITP in cells expressing RADAR systems, we next tested whether RdrB could directly convert ATP to ITP *in vitro*. HPLC analysis demonstrates that purified *E. coli* RdrB is alone sufficient to catalyze rapid conversion of ATP to ITP (Figure 6B). We tested a panel of mononucleotide substrates and observed that *E. coli* RdrB robustly deaminates both ATP and dATP substrates with equal efficiency, while no significant activity was observed with AMP or adenosine mononucleotides (Figure 6C; Figure S6B,C). Mutations to the *E. coli* RdrB conserved histidine tetrad shared with canonical adenosine deaminase enzymes disrupted all detectable deaminase activity (Figure 6D). Finally, we tested a model dsRNA hairpin substrate reported to be RADAR targets *in vivo* (Gao et al., 2020), and observed that under the same reaction condition sufficient for complete conversion of mononucleotides that no RNA A-to-I base-editing occurred (Figure 6E), suggesting that previous observations of RNA editing associated with RADAR defense *in vivo* may be explained by incorporation of inosine mononucleotides into newly synthesized RNA. These data demonstrate that rapid deamination of adenosine nucleotides is a key mediator of RADAR anti-phage defense.

## Discussion

Our results reveal that the RADAR defense system forms a supramolecular complex that targets adenosine nucleotides and protects bacteria from phage replication. Cryo-EM structures define the mechanism of RADAR complex assembly and demonstrate that heptameric RdrA subunits dock around a core dodecameric RdrB shell to create a giant, flower-shaped defense complex (Figure 4). RdrA subunits form a funnel over the RdrB active site, and we show that the RdrB catalytic center is structurally homologous to adenosine deaminases that act on mononucleotide substrates like AMP. Upon phage infection, RADAR induces rapid accumulation of ITP and dITP. Purified RdrB catalyzes conversion of ATP to ITP and dATP to dITP *in vitro*, confirming the direct ability of RADAR to target mononucleotide substrates (Figure 6). In contrast to efficient targeting of adenosine mononucleotides, no significant RNA editing is observed during phage defense or *in vitro* under conditions where ATP is robustly converted to ITP. Rather than deamination of adenosines within RNA, we propose a model in which rapid accumulation of inosine derivatives poisons the nucleotide pool to inhibition of phage replication and abortive infection via cell death (Figure 6G). In agreement with the hypothesis that accumulation of inosine derivatives is poisonous to the cell, overexpression of RdrB in the absence of phage infection results in ITP accumulation and cellular toxicity (Figure S6D, E).

The cellular nucleotide pool has emerged as a common target for host-directed immune responses that disrupt viral replication. In animal cells, SAMHD1 is a triphosphohydrolase enzyme that depletes dNTPs to limit replication of viruses including retroviruses and herpesviruses (Coggins et al., 2020). Recently, nucleotide depletion was identified as a key effector mechanism in anti-phage defense (Hsueh et al., 2022; Tal et al., 2022) and interbacterial competition (Ahmad et al., 2019). In addition to depleting available nucleotides in the cell, immune systems in animal cells and bacteria use viperin enzymes to synthesize nucleotide analogs including CTP derivatives that function as chain-terminators to disrupt replication of diverse viruses and phages (Bernheim et al., 2021; Gizzi et al., 2018). RADAR synthesis of inosine nucleotides provides a mechanistically similar form a defense and limits phage replication by likely increasing nucleotide misincorporation and potentially inhibiting abundant viral enzymes that require ATP hydrolysis for function like phage terminase proteins required for genome translocation and capsid packaging (Rao and Feiss, 2008). The enormous metabolic demand required for genome synthesis creates an opportunity for the host to target free nucleotides and disrupt viral replication. We speculate that an advantage of RADAR synthesis of inosine nucleotides is the relatively low toxicity associated with spurious, low-level ATP-to-ITP conversion in the absence of infection and high antiviral inhibitory activity of elevated levels of inosine synthesis during phage replication and full system activation.

One of the most surprising findings from our structural analysis is that the RADAR components form a giant, highly unusual, multimeric assembly. While some CRISPR immune systems form large complexes to detect invading viral nucleic acid and mount defense (Wang et al., 2022), the specific role of extensive multimerization of repeating subunits in RADAR defense is unknown. However, mutations that disrupt RdrA and RdrB multimerization or RdrA-RdrB complex formation inhibit defense *in vivo* and clearly demonstrate that full assembly is required for inhibition of phage replication (Figure 4). A common theme in mammalian innate immunity, including in inflammasome, toll-like receptor, RIG-I like receptor, and cGAS-STING signaling pathways, is supramolecular complex assembly as a critical step that enhances pathogen detection and downstream immune activation (Kagan et al., 2014). Protein oligomerization and multimerization is also required for effector activation in CBASS immunity(Duncan-Lowey et al., 2021; Hogrel et al., 2022; Morehouse et al., 2022), suggesting that RADAR complex assembly may specifically enhance enzymatic activity to enable anti-phage defense. Building on the new structures of RdrA and RdrB and discovery of mononucleotide targeting, future studies will explain how RADAR complex formation regulates enzymatic function and anti-phage defense.

### Limitations of the Study

Like many recently described antiphage defense systems, it is unclear what specific molecular feature of phage infection induces activation of the RADAR system. We demonstrate that RADAR is rapidly activated upon infection, with ITP and dITP accumulating in the cell within 5 min of early phage replication. We hypothesize that the uncharacterized C-terminal domain of RdrA may act as a sensor, which upon detection of phage replication can promote RADAR complex formation and RdrB adenosine deaminase activity. A major focus of future experiments will be to understand what features of phage infection induce defense and to define how RdrA converts this pathogen recognition event into RADAR complex activation.

## Acknowledgements

The authors thank members of the Kranzusch and Sorek laboratories for helpful discussions. We thank W. Shih’s laboratory for training and use of the JEOL-1400 electron microscope, the Harvard Center for Cryo-Electron Microscopy (HC2EM), the HMS Electron Microscopy Facility, M. Eck for sharing computational resources, and the SBGrid consortium for computational support. This study was supported by the Pew Biomedical Scholars Program (P.J.K.), the Burroughs Wellcome Fund PATH award (P.J.K.), the Mathers Foundation (P.J.K.), the Parker Institute for Cancer Immunotherapy (P.J.K.), European Research Council grant ERC-CoG 681203 (R.S.), Israel Science Foundation grant ISF 296/ 21 (R.S.), the Ernest and Bonnie Beutler Research Program of Excellence in Genomic Medicine (R.S.), the Minerva Foundation and Federal German Ministry for Education and Research (R.S.), the Knell Family Center for Microbiology (R.S.), the Yotam project and the Weizmann Institute Sustainability and Energy Research Initiative (R.S.), and the Dr. Barry Sherman Institute for Medicinal Chemistry (R.S.). B.D.-L. was supported by a Herchel Smith Graduate Research Fellowship, A.G.J. by a Life Science Research Foundation postdoctoral fellowship of the Open Philanthropy Project, and A.M. by a fellowship from the Ariane de Rothschild Women Doctoral Program and, in part, by the Israeli Council for Higher Education via the Weizmann Data Science Research Center.

## Author Contributions

Experiments were designed by B.D.-L., N.T., R.S. and P.J.K. Protein purification, biochemical experiments, and molecular modeling were performed by B.D.-L, with assistance from P.J.K. Phage challenge assays were performed by N.T., T.F. and S.M. Cryo-EM data were collected by A.G.J. and M.M., with assistance from S.R. Cryo-EM data processing was performed by S.R. Cell lysate extraction and analysis of MS data was performed by N.T. Genome analyses were perform by S.D., A.K., and G.A. RNA-seq was performed by N.T., and RNA-seq data analysis was performed by A.M. and N.T. The manuscript was written by B.D.-L., N.T., R.S. and P.J.K., and all authors contributed to editing the manuscript and support the conclusions.

---

**Figure S1.**
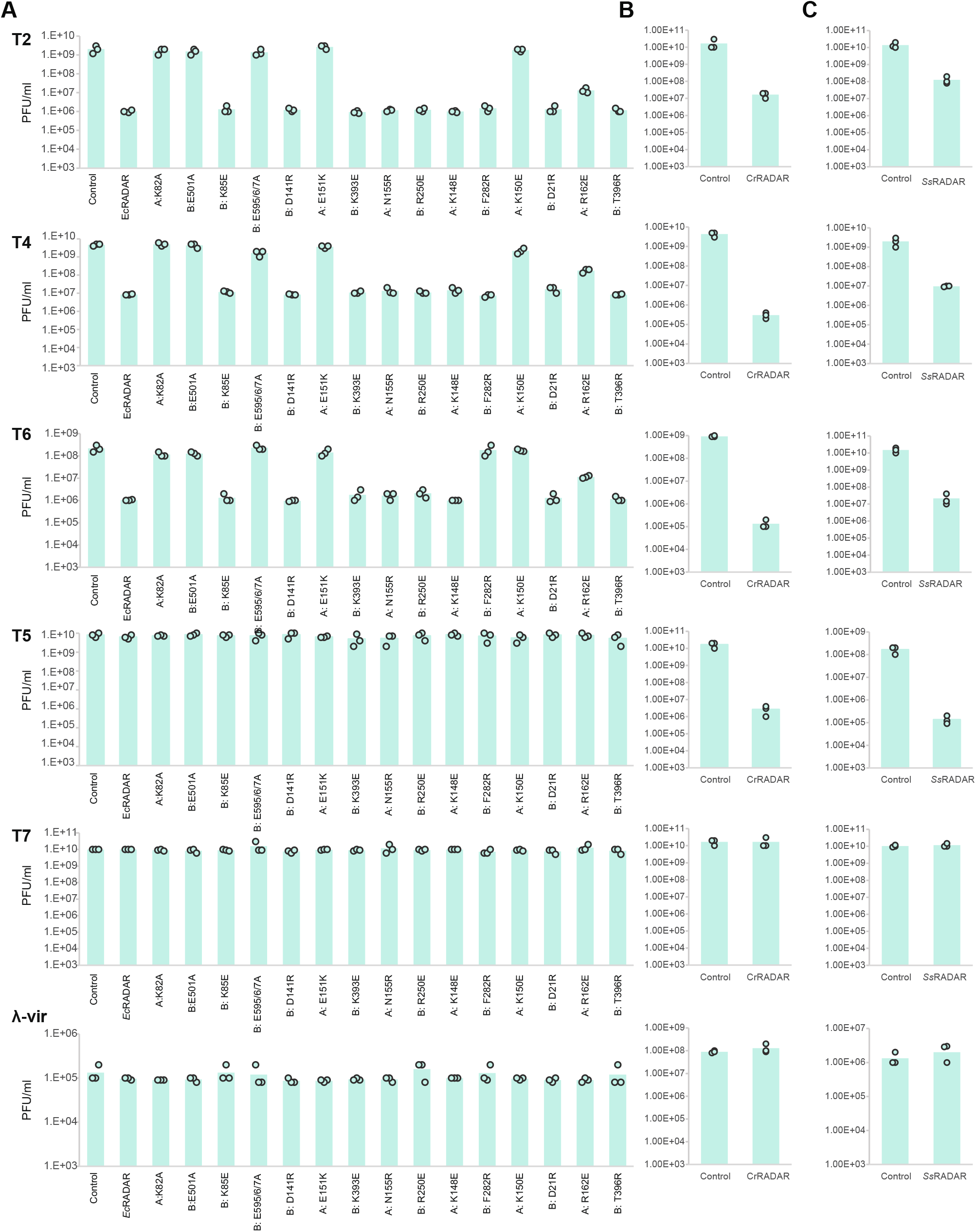
RADAR system from *E. coli* P0304799.3, *C. rodentium* DBS100, and *S. suis* SS993 protect against phage infection, related to Figure 1. (A) Bacteria expressing WT RADAR system from *E. coli* P0304799.3, as well as the system with point mutations in rdrA (‘A’) or rdrB (‘B’), and a negative control that contains an empty vector, were grown on agar plates in room temperature. Tenfold serial dilutions of the phage lysate were dropped on the plates. Data represent plaque-forming units per milliliter for phages tested in this study. Each bar graph represents average of three replicates, with individual data points overlaid. The data for T2-infected WT are also presented in Figures 1E, 3G, and 4G. (B, C) Bacteria expressing RADAR systems from *C. rodentium* DBS100 (B) and *S. suis* SS993 (C) and a negative control that contains an empty vector or a vector expressing GFP (as detailed in the methods section), were grown on agar plates in room temperature. Tenfold serial dilutions of the phage lysate were dropped on the plates. Data represent plaque-forming units per milliliter for phages tested in this study. Each bar graph represents average of three replicates, with individual data points overlaid.

**Figure S2.**
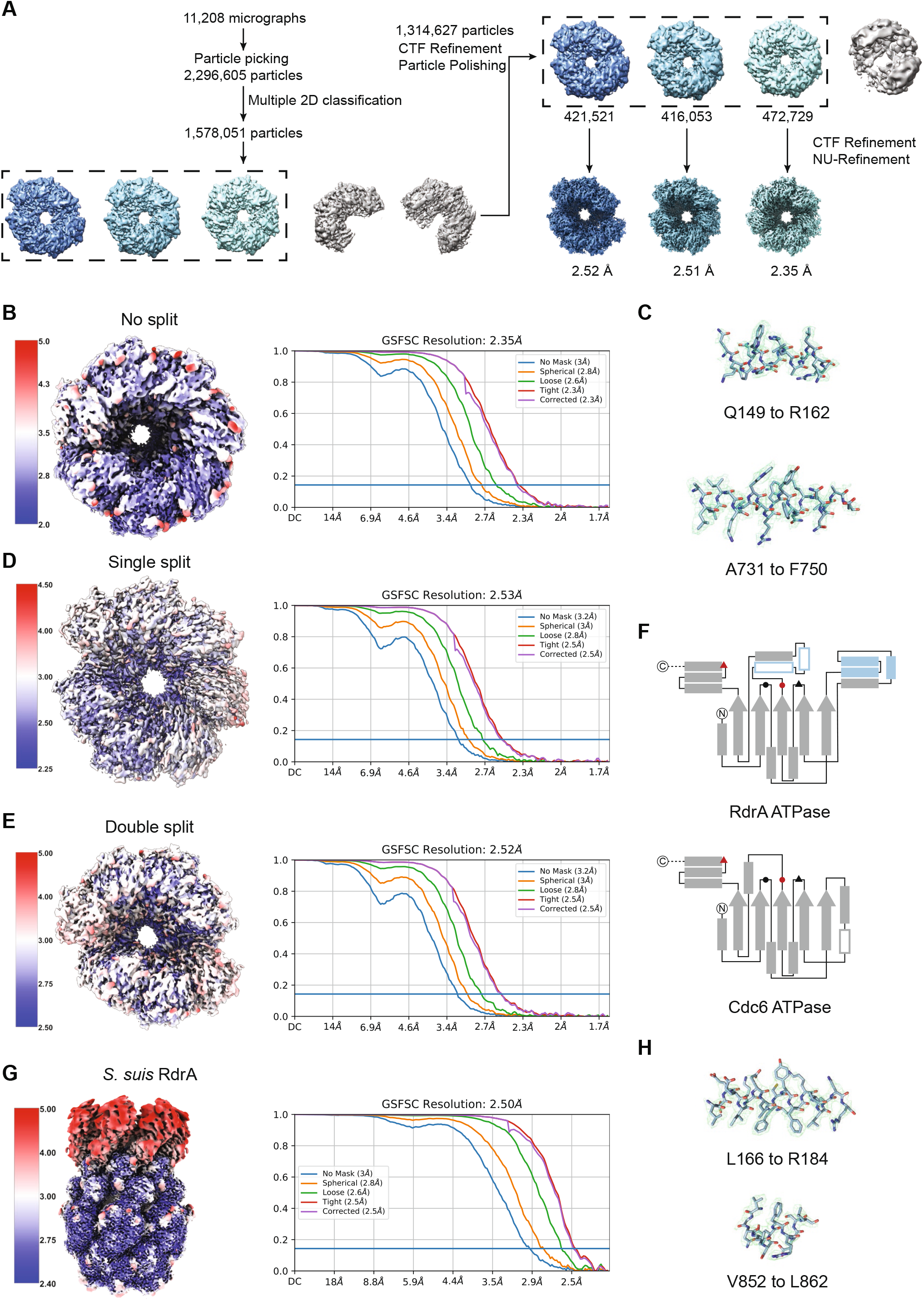
Structure of E. coli and S. suis RdrA ATPase, related to Figure 2. (A) Particle picking and classification strategy for *E. coli* RdrA (B) Local resolution (left) and resolution by Fourier Shell Correlation (FSC) for unsplit *E. coli* RdrA. (C) Example model to map fit from no-split *E. coli* RdrA with the map contoured at 4.0 σ. (D, E) Local resolution (left) and resolution by FSC for *E. coli* RdrA with a single split (D) or a double split (E) (F) Cartoons comparing the ATPase domain of *E. coli* RdrA and Drosophila Cdc6. Conserved ATPase features are annotated as follows: Walker A (black circle), Walker B (black triangle), sensor 1 (red circle), sensor 2 (red triangle). Insertion 1, which forms the “crown”, is shown as filled blue rectangles and insertion 2 is shown as empty rectangles outlined in blue. (G) Local resolution (left) and resolution by FSC for S. suis RdrA. (H) Example model-to-map fit from *S. suis* RdrA with the map contoured to 5.0 σ.

**Figure S3.**
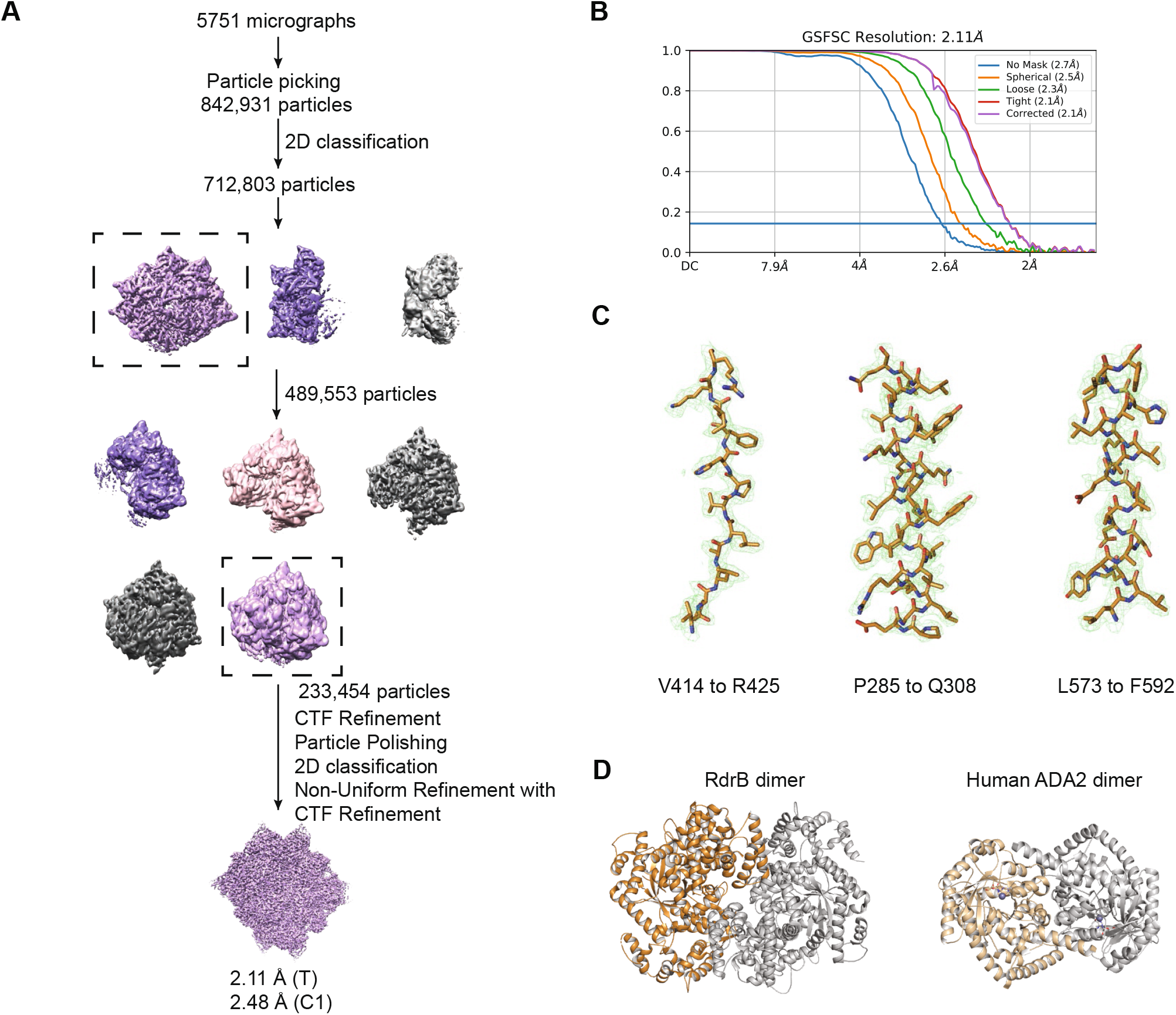
Structure of E. coli RdrA adenosine deaminase, related to Figure 3. (A) Particle picking and classification strategy for E. coli RdrB (B) Resolution by FSC for *E. coli* RdrB. (C) Example model-to-map fit from *E. coli* RdrB with the map contoured to 5.0 σ (D) Comparison of the dimer formed by RdrB along the two-fold interface and the dimer formed by human ADA2

**Figure S4.**
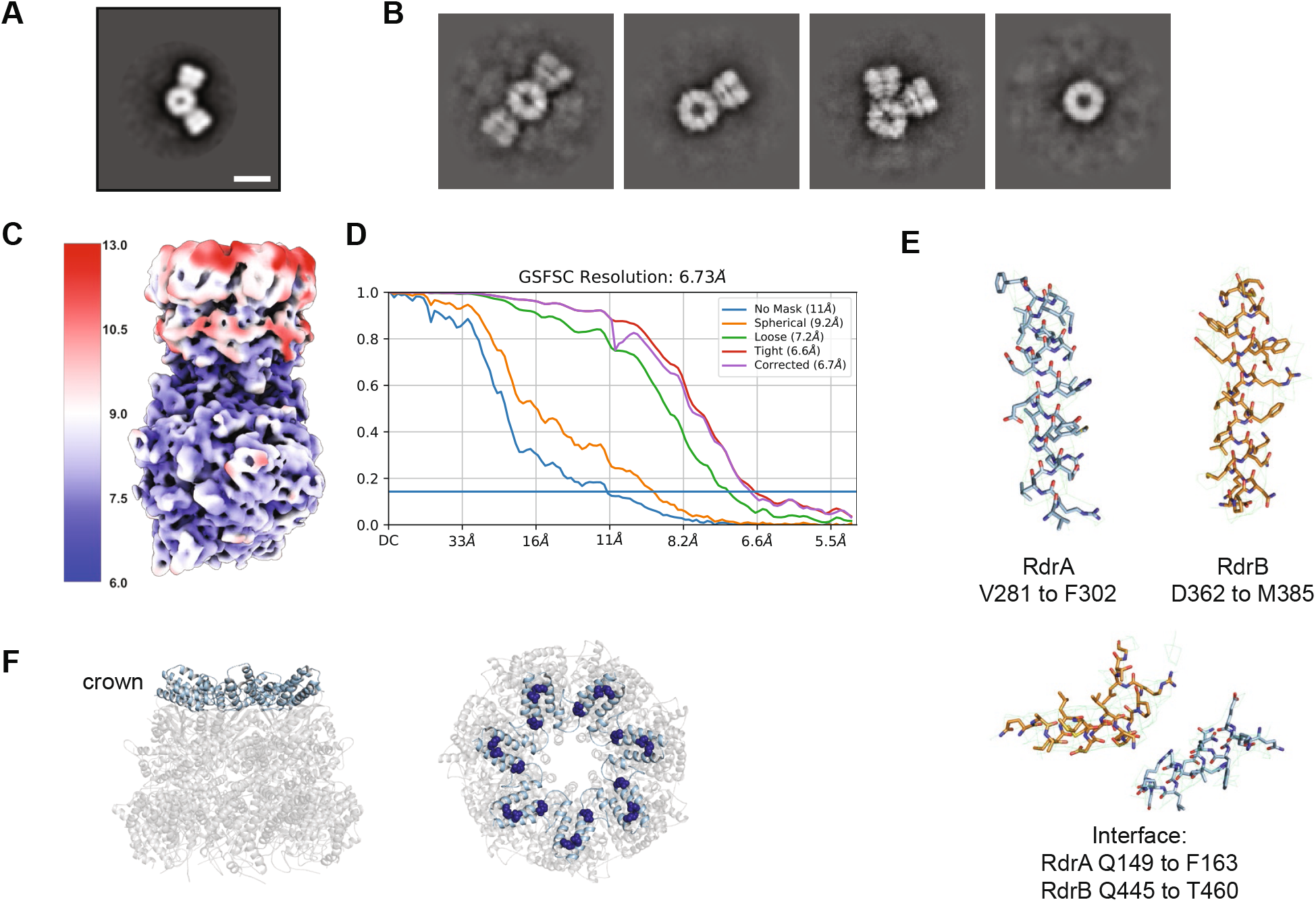
Structure of E. coli RdrA-RdrB supramolecular complex, related to Figure 4. (A) 2D class average from negative stain electron microscopy of RdrA-RdrB complex with a central protein surrounded by two “petals”. Scale bar represents 100 Å. (B) Additional 2D class averages of cryo-EM RdrA-RdrB mixture showing many arrangements with different numbers of RdrA “petals” surrounding a RdrB core. (C, D) Local resolution (C) and resolution by FSC (D) for the *E. coli* RdrA-RdrB complex. (E) Example model to map fit from the *E. coli* RdrA-RdrB complex with the map contoured to 5.0 σ. (F) Cartoon of *E. coli* RdrA heptamer highlighting the “crown” in light blue (left), and the residues mutated to disrupt interface with RdrB (K150E, E151K, R162E as dark blue spheres, right).

**Figure S5.**
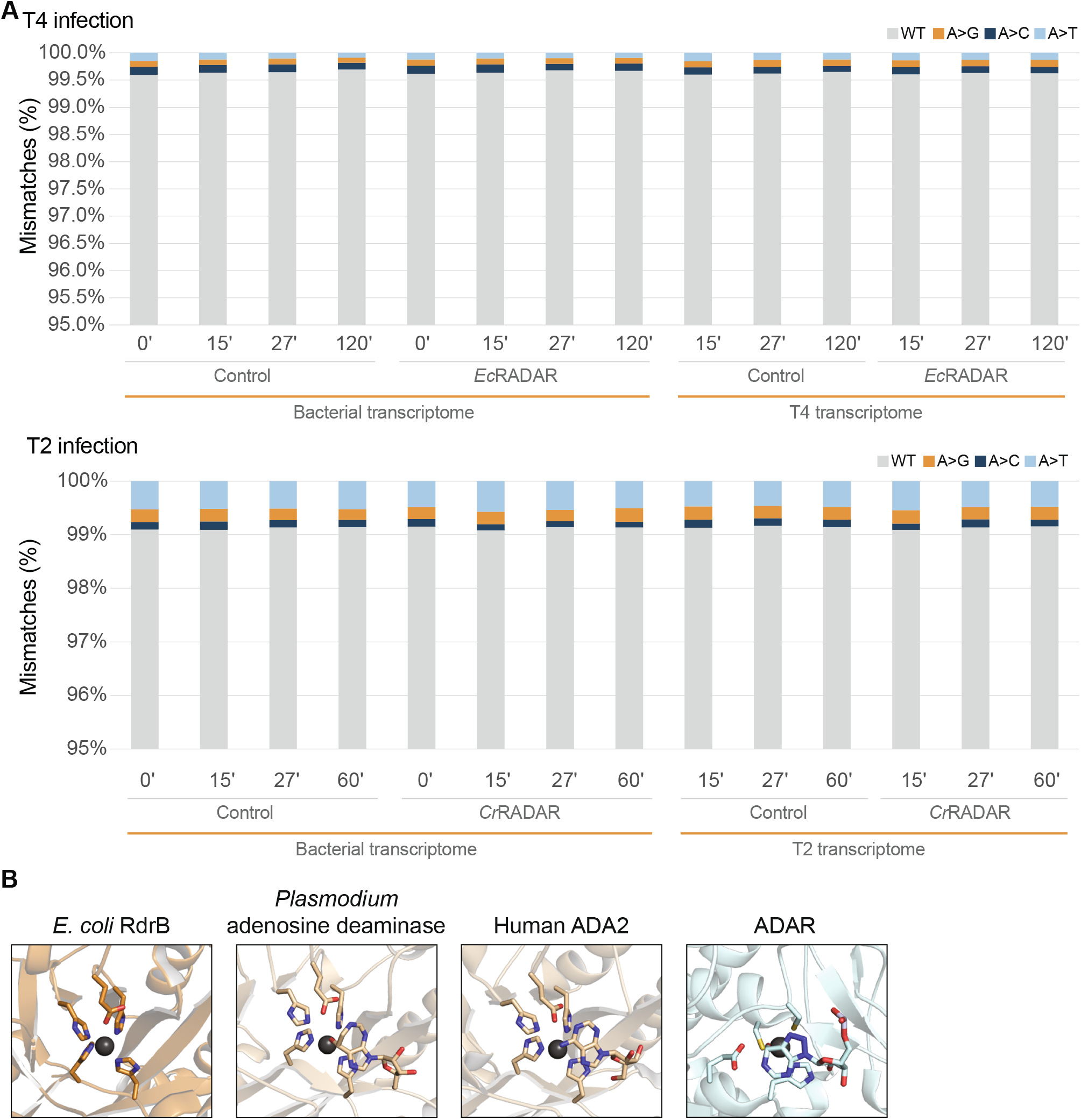
Analysis of deaminase activity of RdrB, related to Figure 5. (A) Mismatches between sequenced RNA and genomic DNA in adenine positions. RNA, extracted from *Ec*RADAR and *Cr*RADAR-expressing cells or control cells infected by phages T4 or T2 (respectively), was sequenced and the resulting reads were aligned to the reference sequences of the *E. coli* host and the respective phage genomes. Shown in the rate of mismatches in all expressed adenosine positions mapped to non-rRNA genes. X-axis depicts the time from the onset of infection, with t=0 reflecting uninfected cells; y-axis is the observed rate of mismatches. (B) Comparison of adenosine deaminase active sites of enzymes that modify monomeric substrates (RdrB, ADA, ADA2) and ADAR, which modifies RNA substrates using a distinct active site and is part of the cytidine deaminase superfamily.

**Figure S6.**
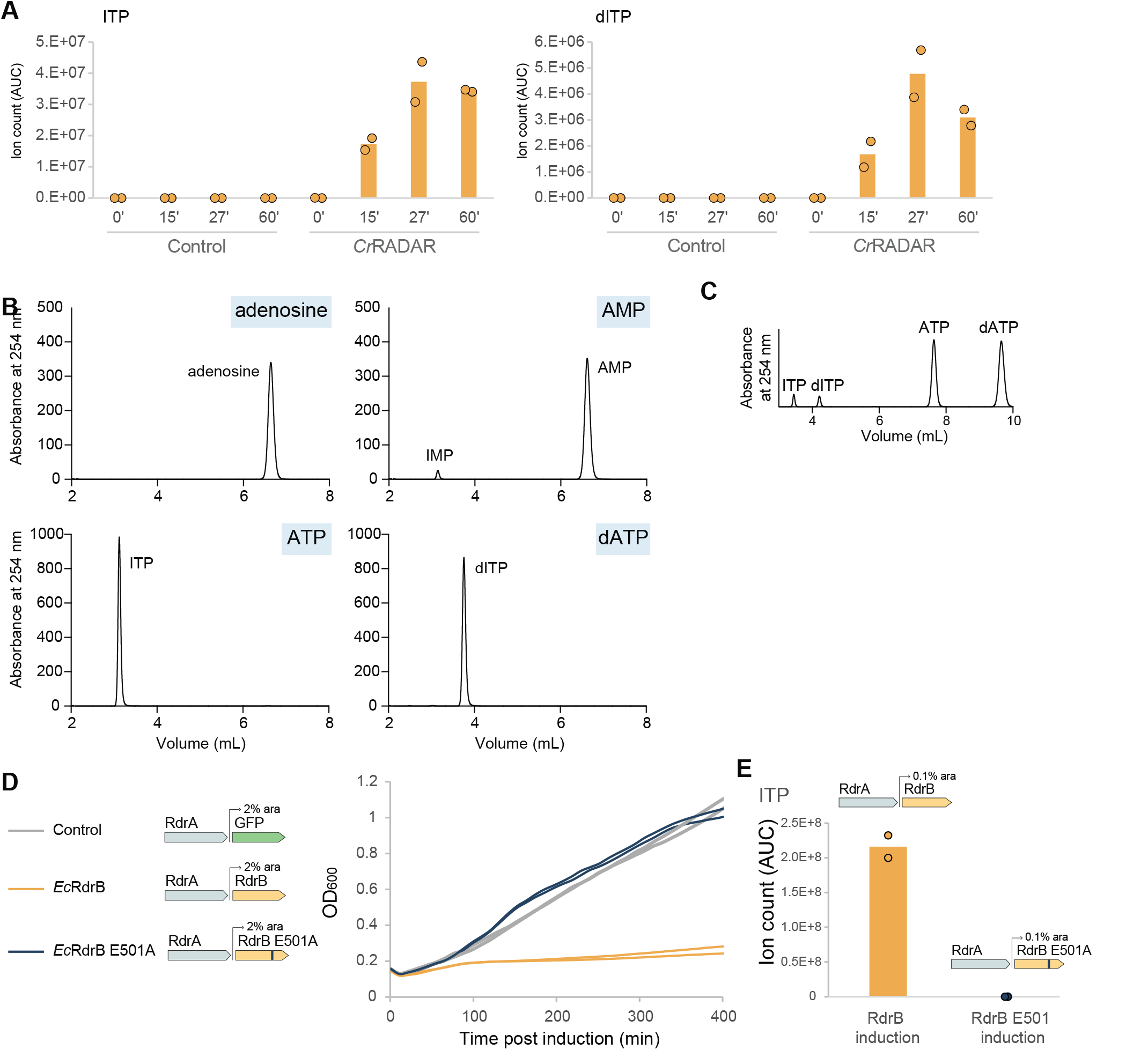
RdrB converts (d)ATP to (d)ITP in vivo and in vitro, related to Figure 6. (A) Ion count (area under curve) of ITP or dITP (respectively) in lysates extracted from WT *Cr*RADAR containing cells, or control cells, as measured by LC-MS/MS. The x axis represents min after infection, with zero representing non-infected cells. Cells were infected by phage T2 at an MOI of 2 at 37°C. Bar graphs represent the average of two biological replicates, with individual data points overlaid. (B) Full HPLC traces of data shown in Figure 6C (C) HPLC analysis of ATP and dATP co-incubated with RdrB, showing similar deamination of both substrates. (D) Growth curves of cells expressing *Ec*RdrB (orange), the mutant *Ec*RdrB E501A (blue) and control cells expressing GFP (grey) in the presence of 2% arabinose without phage infection. Results of two experiments are presented as individual curves.

**Supplementary Table 1. RADAR defense system distribution**

**Supplementary Table 2. Structure statistics**

**Supplementary Table 3. RNA-sequencing**

**Supplementary Table 4. Sequences synthesized in this study**

**Supplementary Table 5. Primers used in this study**

## STAR METHODS

### RESOURCE AVAILABILITY

#### Lead Contact

Further information and requests for resources and reagents should be directed to and will be fulfilled by the Lead Contact, Philip Kranzusch.

#### Materials Availability

This study did not generate new unique reagents.

#### Data and Code Availability

Coordinates of the structures *E. coli* RdrA, RdrB and RdrA-RdrB complex and *S. suis* RdrA have been deposited under the following accession numbers: XXXX.

### EXPERIMENTAL MODEL AND SUBJECT DETAILS

#### Bacterial strains and phages

*E. coli* strain MG1655 (ATCC 47076) was grown in MMB (LB + 0.1 mM MnCl2 + 5 mM MgCl2, with or without 0.5% agar) at 37 °C or room temperature (RT). Whenever applicable, media were supplemented with ampicillin (100 μg/ml), to ensure the maintenance of plasmids. Infection was performed in MMB media at 37°C or RT as detailed in each section. Phages used in this study are listed in the Key Resources Table.

### METHOD DETAILS

#### Protein expression and purification

Recombinant *E. coli* RdrA and RdrB and *S. suis* RdrA were purified using methods previously described (Duncan-Lowey et al., 2021). Briefly, RdrA and RdrB were cloned into an N-terminal 6×His-SUMO2-tagged pET vector and transformed into BL21-RIL *E. coli* (Agilent). Large scale cultures (2–4 liters) were grown for ~5 h at 37°C, then induced with IPTG overnight at 16°C. Bacterial pellets were resuspended and sonicated in lysis buffer (20 mM HEPES-KOH pH 7.5, 400 mM NaCl, 30 mM imidazole, 10% glycerol and 1 mM DTT) and purified using Ni-NTA resin (Qiagen). Ni-NTA resin was washed with lysis buffer supplemented to 1 M NaCl and eluted with lysis buffer supplemented to 300 mM imidazole. The Ni-NTA elution fraction was dialyzed into 20 mM HEPES-KOH pH 7.5, 250 mM KCl, 1 mM DTT overnight while removing the SUMO2 tag with recombinant human SENP2 protease (D364–L589, M497A). RdrA was bound to a Q column (Cytvia) and eluted with a gradient of KCl from 150 mM to 1 M. RdrA and RdrB were each concentrated using a 30K-cutoff concentrator (Millipore) and purified by size exclusion chromatography on a 16/60 Sephacryl 300 column. Proteins were concentrated to >10 mg mL^−1^, flash frozen with liquid nitrogen, and stored at −80°C.

#### Plasmid and strain construction

RADAR operons used for phage challenge assays in this study were synthesized by Genscript Corp. and cloned into the p15a-origin-containing pSG1 plasmid with their native promoters, or into the pBAD plasmid (Thermofisher, cat. #43001), as previously described (Bernheim et al., 2021). Mutants of the system were also synthesized and cloned by Genscript. All synthesized sequences are presented in Table S4. Inducible mutants of RdrB were constructed using Q5 Site directed Mutagenesis kit (NEB, cat. #E0554S), using primers described in Table S5.

#### Plaque assays

Phages were propagated by picking a single phage plaque into a liquid culture of *E. coli* MG1655 grown at 37°C to OD_600_ of 0.3 in MMB medium until culture collapse. The culture was then centrifuged for 10 min at 15,000 × g and the supernatant was filtered through a 0.2 μm filter to get rid of remaining bacteria and bacterial debris. Lysate titer was determined using the small drop plaque assay method as described before (Mazzocco et al., 2009).

Plaque assays were performed as previously described (Mazzocco et al., 2009). Bacteria (*E. coli* MG1655 with pSG1 or pBAD plasmid) and negative control (*E. coli* MG1655 with empty pSG1 or pBad-GFP) were grown overnight at 37°C. Then 300 μL of the bacterial culture was mixed with 30 mL melted MMB agar (LB + 0.1 mM MnCl2 + 5 mM MgCl2 + 0.5% agar, with or without 0.2% arabinose) and left to dry for 1 h at room temperature. 10-fold serial dilutions in MMB were performed for each of the tested phages and 10 μL drops were put on the bacterial layer. Plates were incubated overnight at RT. Plaque forming units (PFUs) were determined by counting the derived plaques after overnight incubation.

#### Liquid toxicity assay

Overnight cultures of bacteria harboring a pSG1 plasmid with *Ec*RdrA and an inducible plasmid (pBbE8k) with different versions of *Ec*RdrB or a GFP control were diluted 1:100 in MMB medium. Cells were incubated at 37°C while shaking at 200 rpm for 1 h. 180 μL of the bacterial culture were transferred into wells in a 96-well plate supplemented with 2% arabinose and incubated at 37°C with shaking in a TECAN Infinite200 plate reader. OD_600_ was followed with measurement every 10 min.

#### Cryo-Electron Microscopy Data Collection

Solutions of purified RdrA, RdrB, and a combination of both, were applied to glow discharged grids and vitrified in liquid ethane. For RdrA and RdrB, concentrations of 2.40 and 2.35 mg/mL were used respectively. For the combination of both, RdrA and RdrB were mixed in a 1:1 ration resulting in final concentrations of 1mg/mL and 0.98 mg/mL respectively. The ideal concentrations to use were based off the preliminary negative stain data. To vitrify the sample on TEM grids, a Mark IV Vitrobot (ThermoFisher) was used. A 3 μL solution of each sample were independently deposited onto 1.2/1.3 Au Quantifoil grid with Carbon mesh then blotted with filter paper for 6 seconds, using a double-sided blot with a force of 5, in a 100% relative humidity chamber at 4°C.

*S.Suis* RdrA, as well as *E. coli* RdrA and RdrB combination grids, were screened and imaged using a Talos Arctica (ThermoFisher) microscope operating at 200 kV and equipped with K3 direct electron detector (Gatan). Approximately 300 and 240 movies were acquired of the RdrA and the RdrA and RdrB combination sample, respectively, using SerialEM software version 3.8.6 at a pixel size of 1.1 Å, a total dose of 42.02 e− /Å^2^, dose per frame of 1.05 e− /Å^2^. A defocus range of −0.7 to −2.0 μm was used for *E. coli* RdrA and RdrB combination grids while a defocus range of −0.5 to −3.0 μm was used for *S.Suis* RdrA.

*E.Coli* RdrA and *E.Coli* RdrB grids were screened and imaged using a Titan Krios microscope operating at 300 kV and equipped with a K3 direct electron detector with energy filter (Gatan). All data was acquired using SerialEM software version 3.8.6 at a pixel size of 0.825 Å and a defocus range of −0.5 to −2.5 um. Approximately 11,200 movies of *E.Coli* RdrA were collected at a total dose of 48.8 e− /Å^2^, dose per frame of 1.16 e− /Å^2^. Approximately 5,700 movies of *E.Coli* RdrB were collected with a total dose of 48.8 e− /Å^2^, dose per frame of 1.16 e− /Å^2^.

#### Cryo-EM Data Processing

Dose-fractionated images of *E. coli* RdrA were gain normalized and motion corrected with MotionCor2 (v1.3.1) (Zheng et al., 2017) followed by CTF and defocus value determination in CTFFIND4 (Rohou and Grigorieff, 2015). Particle picking was carried out in crYOLO (Wagner et al., 2019) resulting in 2,296,605 initial particles. Following multiple rounds of 2D classification in RELION (Scheres, 2012) to remove erroneous picks, contamination, and “junk” particles 1,578,051 particles representing intact RdrA were obtained. Heterogeneous refinement in cryosparc (Punjani et al., 2017) identified three different populations of RdrA, totaling 1,314,627 particles. Following particle polishing (Zivanov et al., 2019) and CTF refinement (Zivanov et al., 2020)in RELION on the combined data a further cycle of heterogeneous refinement followed by Non-uniform refinement (Punjani et al., 2020) and additional CTF refinement in cryosparc resulted in the “no split”, “single split”, and “double split” reconstructions at resolutions of 2.3, 2,5 and 2.5Å respectively (Figure S2A).

*E. coli* RdrB was processed in a similar manner with 842,931 particles initial identified, leading to 712,803 after 2D classification. Following multiple rounds of heterogeneous refinement in cryopsarc 233,454 particles were subjected to polishing, CTF refinement, particle polishing, additional 2D classification and finally Non-uniform refinement resulting in a 2.1Å reconstruction with T symmetry (2.5Å C1) (Figure S3A).

The small set of *E. coli* RdrA-RdrB complex images were motion corrected using the RELION implementation, followed by CTFFIND4 and particle picking in crYOLO, resulting in 20,409 particles. All particles were subjected to ab initio reconstruction into three classes in cryosparc. 9,236 particles were identified corresponding to the complex, resulting in a 6.7Å reconstruction following refinement.

Dose-fractionated images of *S. suis* RdrA were gain normalized and motion corrected with MotionCor2 (v1.4.0) followed by CTF and defocus value determination in CTFFIND4 (Rohou and Grigorieff, 2015). crYOLO models were trained to identify potential “monomer” (single heptametic ring) and “dimer” (double stacked rings) species. These were then combined and duplicate particles removed prior to 2D classification in RELION which resulted in 573,499 particles after “junk” removal. These particles then underwent multiple rounds of heterogeneous refinement within cryosparc alongside CTF refinement resulting in 193,305 particles in total. These particles were polished in RELION before a final round of CTF refinement and Non-uniform refinement in cryosparc resulting in the final C7 reconstruction at 2.5Å (2.7 Å C1).

Structural biology applications other than cryosparc used in this project were compiled and configured by SBGrid (Morin et al., 2013).

#### Negative Stain Electron Microscopy

*E coli* RdrA and RdrB were mixed to a final concentration of 50 nM for each protein in buffer containing 100 mM KCl, 50 mM HEPES pH 7.5, and 1 mM TCEP. 4 μL samples were applied to glow-discharged copper grids (Electron Microscopy Sciences, cat. #FCF400-Cu), stained with 2% uranyl formate, and imaged on a JEOL-1400 at 80kV. To determine sample quality for subsequent cryo-EM analysis, 30 micrographs were acquired at 40kx magnification (3.3 pixels nm^−1^) and used for 2D classification in RELION (Scheres 2012 PMID 23000701). Select 2D classes representing different arrangements of RdrA petals around the RdrB core were analysed in ImageJ to add scale bars.

#### RNA sequencing

Overnight cultures of bacteria (*E. coli* MG1655 harboring pSG1-*Ec*RADAR or pSG1-*Cr*RADAR plasmid) or negative control (*E. coli* MG1655 with the pSG1 plasmid) were diluted 1:100 in 60 mL of MMB medium and incubated at 37°C while shaking at 200 rpm until early log phase (OD600 of 0.3). 10 mL samples of each bacterial culture were taken and centrifuged at 4000 rpm for 5 min at 4°C. The pellets were flash frozen using dry ice and ethanol. The remaining cultures were infected by phage T4 or T2, at a final MOI of 2. 10 mL samples were taken throughout infection at 0, 15, 27 and 120 min post infection (for *Ec*RADAR), or 0, 15, 27 and 60 min post infection (for *Cr*RADAR), and centrifuged and flash frozen as described above. RNA extraction was performed as described previously (Dar et al., 2016). Briefly, frozen pellets were re-suspended in 1 mL of RNA protect solution (FastPrep) and lysed by Fastprep homogenizer (MP Biomedicals). RNA was extracted using the FastRNA PRO blue kit (MP Biomedicals, 116025050) according to the manufacturer’s instructions. DNase treatment was performed using the Turbo DNA free kit (Life Technologies, AM2238). RNA was subsequently fragmented using fragmentation buffer (Ambion-Invitrogen, cat. #10136824) at 72°C for 1 min and 45 s. The reactions were cleaned by adding ×2.5 SPRI beads (Agencourt AMPure XP, Beckman-Coulter, A63881). The beads were washed twice with 80% ethanol and air dried for 5 min. The RNA was eluted using water. Ribosomal RNA was depleted by using the Ribo-Zero rRNA Removal Kit (epicentre, MRZB12424). Strand-specific RNA-seq was performed using the NEBNext Ultra Directional RNA Library Prep Kit (NEB, E7420) with the following adjustments: all cleanup stages were performed using ×1.8 SPRI beads, and only one cleanup step was performed after the end repair step. Following sequencing on an Illumina NextSeq500, sequenced reads were demultiplexed and adapters were trimmed using ‘fastx clipper’ software with default parameters. Reads were mapped to the bacterial and phage genomes by using NovoAlign (Novocraft) v3.02.02 with default parameters as previously described (Dar et al., 2016). Reads mapped to rRNA genes were discarded. Reads mapping equally well to multiple positions in the reference genome, as well as reads containing insertions and deletions as compared to the reference genome, were also discarded. Only reads mapping to the antisense strand of annotated genes were used for the mutation analyses, as these reads represent cDNA generated from the mRNA. Mutations from reference genomes were identified and quantified by counting each mismatch across the transcriptome. Frequency of mismatches was compared between control and RADAR samples throughout the infection time course.

#### Cell lysate preparation

Overnight cultures of *E. coli* harboring the defensive system and negative controls were diluted 1:100 in 250 mL MMB medium (with or without 0.2% arabinose, as described in Table S4 and grown at 37°C (250 rpm) until reaching OD600 of 0.3. The cultures were infected by T2 or T4 at a final MOI of 2. Following the addition of phage, at 5, 15 and 60 or 120 min post infection (plus an uninfected control sample), 50 mL samples were taken and centrifuged for 5 min at 15,000 × g. Pellets were flash frozen using dry ice and ethanol. The pellets were re-suspended in 600 μL of 100 mM phosphate buffer at pH 8 and supplemented with 4 mg mL^−1^ lysozyme. The samples were then transferred to a FastPrep Lysing Matrix B 2 mL tube (MP Biomedicals cat. #116911100) and lysed using a FastPrep bead beater for 40 s at 6 m/s (two cycles). Tubes were then centrifuged at 4°C for 15 min at 15,000 × g. Supernatant was transferred to Amicon Ultra-0.5 Centrifugal Filter Unit 3 kDa (Merck Millipore cat. #UFC500396) and centrifuged for 45 min at 4°C at 12,000 × g. Filtrate was taken and used for LC-MS analysis.

#### Detection of inosine compounds in cell lysates by HPLC-MS

Profiling of polar metabolites was done as previously described (Campbell and Yamada, 1989) with minor modifications as described below. In brief, the analysis was performed using an Acquity I class UPLC System combined with a mass spectrometer (Thermo Exactive Plus Orbitrap), which was operated in a positive ionization mode using a mass range of 200–800 m/z. The LC separation was done using the SeQuant Zic-pHilic (150 mm × 2.1 mm) with the SeQuant guard column (20 mm × 2.1 mm) (Merck). The mobile phase B was acetonitrile and mobile phase A was 20 mM ammonium carbonate plus 0.1% ammonia hydroxide in water. The flow rate was kept at 200 μl min^−1^ and the gradient was as follows: 75% B (0–2 min), decrease to 25% B (2–14 min), 25% B (14–18 min), increase back to 75% B (18–19 min), 75% B (19–23 min). Inosine derivatives peaks were identified in the data using MSMS fragmentation, by identifying the inosine base signature as well as phosphates, ribose or deoxyribose. Area under the peak was quantified using MZmine 2.53 (Pluskal et al., 2010) with an accepted deviation of 5 ppm.

#### Analysis of base editing by HPLC

All RdrB reactions were carried out at 37°C in standard reaction conditions: 50 mM KCl, 50 mM HEPES pH 7.5, 1 mM TCEP, 10 mM MgCl_2_, 100 uM ZnSO_4_, and 1 μM *E. coli* RdrB. For monomeric substrates (ATP, dATP, AMP, adenosine), 1 mM final concentration of substrate was used, and reactions were incubated for 30 min before heating to 90°C to end reactions. For comparison of ATP and dATP conversion, reactions containing 1 mM ATP and 1 mM dATP were incubated at 37°C for 10 min. Hairpin RNA experiments contained 1 μM hairpin RNA and were at 37°C for 1 h. Reactions were treated with calf intestinal phosphatase (New England Biolabs) and hairpin RNA samples were concurrently treated with P1 nuclease to release monomeric NMPs for analysis, then samples were spun through a 0.2 μm filter. Analysis was carried out using a C18 column (Agilent Zorbax Bonus-RP 4.6×150 mm, 3.5-micron). The column was heated to 40°C and run at 1 mL min^−1^ with a mobile phase of 50 mM NaH_2_PO_4_ (pH 6.8 with NaOH) supplemented with 3% acetonitrile.

### QUANTIFICATION AND STATISTICAL ANALYSIS

Statistical details for each experiment can be found in the figure legends and outlined in the corresponding methods details section. Bar graphs show the average of replicates with individual points overlaid, unless stated otherwise.

